# Photoperiodic Regulation of SK Channels in Dorsal Raphe Serotonin Neurons

**DOI:** 10.1101/2025.03.29.646123

**Authors:** Manuel A. Giannoni-Guzmán, José C. Zepeda, Anna Kamitakahara, Valerie Magalong, Pat Levitt, Brad A. Grueter, Douglas G. McMahon

## Abstract

Day length, or seasonal photoperiod, shapes mood and affective behaviors but the neural mechanisms underlying these effects are still being defined. Serotonin neurons of the dorsal raphe nucleus (DRN) are critical regulators of affective behaviors and photoperiod modulates their excitability and ongoing activity. Here, we investigated the influence of seasonal photoperiod on the function and expression of small conductance calcium-activated potassium (SK) channels which mediate the afterhyperpolarizing potential (AHP) in dorsal raphe serotonin neurons. Building on previous work demonstrating that photoperiod modulates serotonergic excitability and behavior, we hypothesized that day length influences SK channel activity, thereby contributing to differences in neuronal excitability observed between Long, Equinox, and Short photoperiod conditions. Using multi-electrode array recording of DRN slices we found a significant dose-dependent increase in spike rate to the application of the SK channel inhibitor apamin, indicating that SK channels indeed influence the spike rate of dorsal raphe serotonin neurons. In addition, DRN neurons in slices from Long photoperiod mice exhibited less pronounced responses to apamin relative to those from Short photoperiod mice, suggesting reduced function or expression of SK channels in Long photoperiod. Indeed, whole-cell recordings demonstrated that SK channel–mediated AHP currents were reduced in Long photoperiod mice. However, there were no significant differences in expression levels of the SK3 subunit (Kcnn3) in DRN serotonin neurons across photoperiod conditions as determined by single molecule fluorescence in situ hybridization. Overall, these findings indicate that photoperiod modulates SK channel function in DRN serotonin neurons likely at a post-transcriptional level. This study advances our understanding of how seasonal cues influence intrinsic neuronal properties and provides a mechanistic link between photoperiod, serotonergic excitability, and mood-related behaviors. The identification of SK channels as modulators of photoperiodic effects may offer novel therapeutic targets for mood disorders associated with dysregulated serotonin signaling.

## Introduction

Environmental signals sculpt brain development, resulting in changes in overall behavior and physiology (Kolb & Whishaw, 1998; Meaney, 2001). In mammals, early-life experiences can have lasting effects on neural development, shaping behavioral phenotypes into adulthood (Andersen, 2015; Boulle et al., 2016; Damulewicz et al., 2022; Pyter & Nelson, 2006; Rincón-Cortés & Sullivan, 2014). Day length (photoperiod) is a key environmental cue that organisms use to synchronize their internal biological clocks with seasonal changes, thereby timing critical processes such as reproduction and development to maximize offspring survival (Goldman, 2001; Nakane & Yoshimura, 2019). Work in rodents has shown that exposure to seasonal photoperiods alters the function of serotonergic and dopaminergic neurons, potentially leading to changes in behavior (Green et al., 2015; Jameson et al., 2023, 2024; Pyter & Nelson, 2006; Siemann et al., 2019, 2020). Research in human populations suggests associations between developmental photoperiod and lifetime risk for major depression (Devore et al., 2018; Lewis et al., 2024; Viejo-Romero et al., 2024).

In mice, perinatal photoperiod exposure results in enduring changes to the master circadian clock located in the suprachiasmatic nucleus (SCN) (Ciarleglio et al., 2011; Cox et al., 2024). The dorsal raphe nucleus (DRN), the largest serotonergic nuclei in the brain (Hornung, 2010), receives light input from both the master clock and the circadian visual system (Gooley et al., 2003; Morin, 1999, 2013). Using the C3Hf+/+ mice strain, which possesses intact melatonin signaling, our group has shown that photoperiodic signals influence serotonergic brain circuits (Giannoni-Guzmán et al., 2020; Green et al., 2015; Siemann et al., 2019). Behaviorally, mice raised under short (8:16hr LD) (winter-like) photoperiods present increased depressive and anxiety-like behaviors as adults when compared to mice raised under Long (16:8hr LD) (summer-like) photoperiod (Green et al., 2015; Siemann et al., 2019). Serotonin neurons in DRN slices from Long mice exhibited increased excitability, characterized by depolarization of the resting membrane potential and a decreased afterhyperpolarization (AHP) amplitude (Green et al., 2015). We have shown that the transcriptional expression of the twin-pore K+ channel TREK-1—a key ion channel setting the resting membrane potential and regulating neuronal excitability (Talley et al., 2001) — is significantly decreased in serotonin neurons from Long photoperiod mice compared to those from Short photoperiod mice (Giannoni-Guzmán et al., 2020). The behavioral, transcriptional and neural effects photoperiodic programming were negated in melatonin receptor 1 (MT-1) knockout mice, indicating that melatonin signaling is required for the changes induced by photoperiod in DRN serotonin neurons (Giannoni-Guzmán et al., 2020; Green et al., 2015).

Significant gaps remain in our understanding of the intrinsic ion channel mechanisms that underlie photoperiodic programming of this system, such as changes observed in the AHP of neurons. In serotonin neurons of the DRN, the AHP is carried primarily by small conductance calcium-activated K+ channels (SK) (Stocker, 2004), with the SK3 channel presenting the highest expression in this brain region (Stocker & Pedarzani, 2000). SK channel currents are selectively inhibited by the peptide toxin apamin (IC50 = 0.7–4 nM), which binds to the external pore region of the channel, thereby blocking potassium efflux and attenuating the AHP that regulates neuronal excitability. (Crespi, 2009; Grunnet et al., 2001; Kuzmenkov et al., 2022; van der Staay et al., 1999). Apamin has been shown to reduce depressive-like behavior in rodents when administered systemically as well as modulating dorsal raphe serotonergic neuron firing, highlighting its potential to regulate affective behavior through SK channel inhibition (Sargin et al., 2016).

Here we hypothesize that photoperiod modulates SK channel currents in DRN serotonergic neurons, leading to differential changes in the excitability and activity of DRN serotonin neurons between photoperiods. To test this hypothesis, we first determined how pharmacological inhibition of SK channels affect serotonin neuron activity under different photoperiod conditions using multielectrode array recordings. We then quantified SK currents in DRN serotonin neurons across photoperiods employing patch-clamp electrophysiology. Lastly, we examined whether photoperiod alters *Kcnn3* (SK3) expression in DRN serotonin neurons using RNAscope.

## Materials and Methods

### Animals and Housing

All mice used in our experiments were from the C3Hf+/+ mouse strain (a gift of Gianluca Tosini, Morehouse School of Medicine, Atlanta, GA). These mice (C3A.BLiA-Pde6b+/J; JAX stock #001912) have intact melatonin signaling and do not carry the *rd* allele that causes retinal degeneration in the C3 H/HeJ parent strain (Baba et al., 2009). Mice were paired and placed on either Long (16 hours of light and 8 hours of darkness), Equinox (12 hours of light 12 hours of darkness) or Short (8 hours of light and 16 hours of darkness) photoperiods. Litters were maintained in group housing under the same photoperiod until experiments were performed between postnatal days 40 and 60. First litters from breeders were not used. Tissue harvesting occurred between 11:00am-12:00pm local time, the mid-day point of all light cycles. Male and female animals were used in equal numbers in each cohort. Experiments were performed in accordance with the Vanderbilt University Institutional Animal Care and Use Committee and National Institutes of Health guidelines.

### Multielectrode Array Electrophysiological Recording

Mice were euthanized, brains were extracted and mounted in cold, oxygenated (95% O_2_-5%CO_2_) dissecting media (in mM: 114.5 NaCl, 3.5 KCl, 1 NaH_2_PO4, 1.3 MgSO_4_, 2.5 CaCl, 10 D(+)-glucose, and 35.7 NaCHO_3_), and 300μm thick coronal slices were taken using a Vibroslicer (Campden Instruments). Dorsal raphe nuclei were isolated by removing the extraneous cortical tissue and placed sample in a slice chamber full of room temperature, oxygenated, extracellular recording media (in mM: 124 NaCl, 3.5 KCl, 1 NaH2PO_4_, 1.3 MgSO_4_, 2.5 CaCl_2_,10 D(+)-glucose, and 20 NaHCO3).

Multielectrode array recordings were performed by dissecting out the dorsal raphe nucleus after slicing, as previously described (Giannoni-Guzmán et al., 2020; Manz et al., 2021). The tissue was then placed on a perforated electrode array and immobilized with a harp for recording. To elicit spontaneous firing of serotonin neurons, 40 μM tryptophan and 3 μM phenylephrine were added to the recording solution which was perfused (1.3 ml/sec) over the slice. After 45 minutes of placing the slice in solution recording began. For SK channel inhibition, apamin was perfused (1.3 ml/sec) over the slice in a serial manner at concentrations of 0, 1nM, 5nM, 50nM and 100nM. Each dose was perfused for 5 minutes to ensure the desired concentration in the bath. Lastly, the 5-HT_1_A receptor agonist 8-OH DPAT was perfused into the solution at 5μM concentration to identify serotonergic cells by silencing.

### Whole-cell patch-clamp electrophysiology

Male and female mice were killed, their brains were removed, and 250 μm thick coronal brain slices were collected using a Leica VT1200S Vibratome. Slices were prepared in oxygenated (95% O_2_; 5%CO_2_) ice-cold N-methyl-D-glucamine (NMDG)-based solution (in mM: 2.5 KCl, 20 HEPES, 1.2 NaH2PO4, 25 Glucose, 93 NMDG, 30 NaHCO3, 5.0 sodium ascorbate, 3.0 sodium pyruvate, 10 MgCl_2_, and 0.5 CaCl_2_-2H_2_O) and recovered for 10 min in the same solution at 34 °C. Slices were then recovered for 1 h in oxygenated artificial cerebrospinal fluid (ACSF) containing (in mM: 119 NaCl, 2.5 KCl, 1.3 MgCl_2_-6H_2_O, 2.5 CaCl_2_-2H_2_O, 1.0 NaH_2_PO_4_-H_2_O, 26.2 NaHCO_3_, and 11 glucose; 287–295 mOsm). During experiments, slices were continuously perfused with oxygenated ACSF containing 2 μM phenylephrine to stimulate spontaneous action potentials at a rate of 2 mL/min and a temperature of 30-32 °C. Whole-cell patch-clamp electrophysiology was performed using a CV-7B head stage, Multiclamp 700B Amplifier, Axopatch Digidata 1550 digitizer, and 3-6 MΩ glass recording micropipettes (Sutter P1000 Micropipette Puller). Whole-cell patch clamp recordings were performed using a K+-based intracellular solution (in mM: 135 K+-gluconate, 5 NaCl, 2 MgCl_2_, 10 HEPES, 0.6 EGTA, 3 Na_2_ATP, 0.4 Na_2_GTP; 285-292 mOsm). In order to identify serotonergic cells, spontaneous action potentials were recorded from neurons using the same intracellular solution before and during a bath application of the 5-HT_1_A receptor agonist 8-OH DPAT (5 μM), only cells whose spontaneous action potentials decreased by >50% in the presence of 8-OH DPAT were included in this study. To elicit afterhyperpolarization currents, a 2ms pulse to + 30 mV from the holding potential of – 60 mV was applied in the voltage clamp configuration.

### In situ hybridization quantification

Whole brain tissue was collected (n = 6 per group) and submerged into isopentane for 25 seconds, placed in crushed dry ice for approximately 1 minute, and stored in a 50mL falcon tube at -80°C. 20 μm sections were prepared per animal targeting the dorsal raphe nucleus from -5.5mm to -5.75mm from Bregma. Tissue sections were then processed by single molecule fluorescence in situ hybridization using RNAscope™ (Advanced Cell Diagnostics). Specific probes targeted the transcripts *Kcnn3* (SK3) and *Tph2* with fluorescent dyes Alexa488, and Atto647, respectively. Once tissue was processed, laser scans for each fluorescent channel containing the dorsal raphe nucleus for each brain slice were captured using a Zeiss LSM510 confocal scanning microscope (Carl Zeiss Microscopy Gmbh, Jena, Germany). Sixteen sequential scans at 40x of 512×512 pixels were taken by the microscope and stitched together to produce a high-resolution composite of the area. Laser settings and pinhole sizes were the same for all the captures. Image analysis was performed using the open source software Fiji (https://imagej.net/Fiji). Colocalization of *Kcnn3* with *Tph2* was established using the plugin ComDet (https://github.com/ekatrukha/ComDet) (Fréal et al., 2019). Briefly, the plugin determines small ROIs for each channel independently and determines if they colocalize spatially with up to 4 pixels of distance from each other. Number of colocalized particles is reported as a proxy for gene expression levels.

### Statistical Analysis

Multielectrode array traces were analyzed using offline sorting (Plexon) and spikes were sorted using a combination of manual identification and automatic k-means-based sorting software. Graphs and statistics were prepared using GraphPad PRISM 10. Two-way ANOVA was performed for each of the cohorts using photoperiod and inhibitor dose as independent variables. Since we used male and female mice in each of the cohorts in equal numbers, sex differences were analyzed for all cohorts, however, no significant differences were observed.

## Results

### Photoperiodic Modulation of SK Channel Inhibition on DRN Serotonin Neuron Activity

To examine the role of SK channels on DRN serotonin firing, we inhibited SK channels in a dose-dependent manner using apamin (0, 1 nM, 5 nM, 50 nM, and 100 nM) and quantified the spike rate of DRN serotonin neurons. Multielectrode array recordings of acute brain slices of the dorsomedial region of the DRN of mice (C3Hf+/+) conceived and raised into adulthood in either Long (summer-like), Equinox (spring/fall-like), or Short (winter-like) photoperiods were used. The spontaneous firing of serotonin neurons was identified in each recording by suppression of spike frequency in response to 8-OHDPAT at the end of each recording, as previously described (Giannoni-Guzmán et al., 2020; Green et al., 2015). We hypothesized that in Long photoperiod mice, which present increased firing rate and increased excitability (Green et al., 2015), inhibition of SK channels would have a reduced effect, while in Short photoperiod mice, we would observe an increased response to SK channel inhibition.

We observed a significant dose-dependent increase in the spike rate of Long and Short photoperiod cohorts (Figure 1A). Repeated measures two-way ANOVA revealed significant independent effects for apamin dose (p < 0.0001****) and photoperiod (p < 0.001**). The interaction between photoperiod and apamin dose was also significant (p < 0.05*), with the most pronounced effect observed in the Short photoperiod cohort. Tukey’s multiple comparisons test revealed a significantly stronger effect of apamin on serotonin neurons from the Short photoperiod cohort (adjusted p = 0.0287, comparing 0 vs. 100 nM). This result suggests that the SK channel currents in dorsal raphe neurons are under the influence of photoperiodic programming.

**Figure 1.**
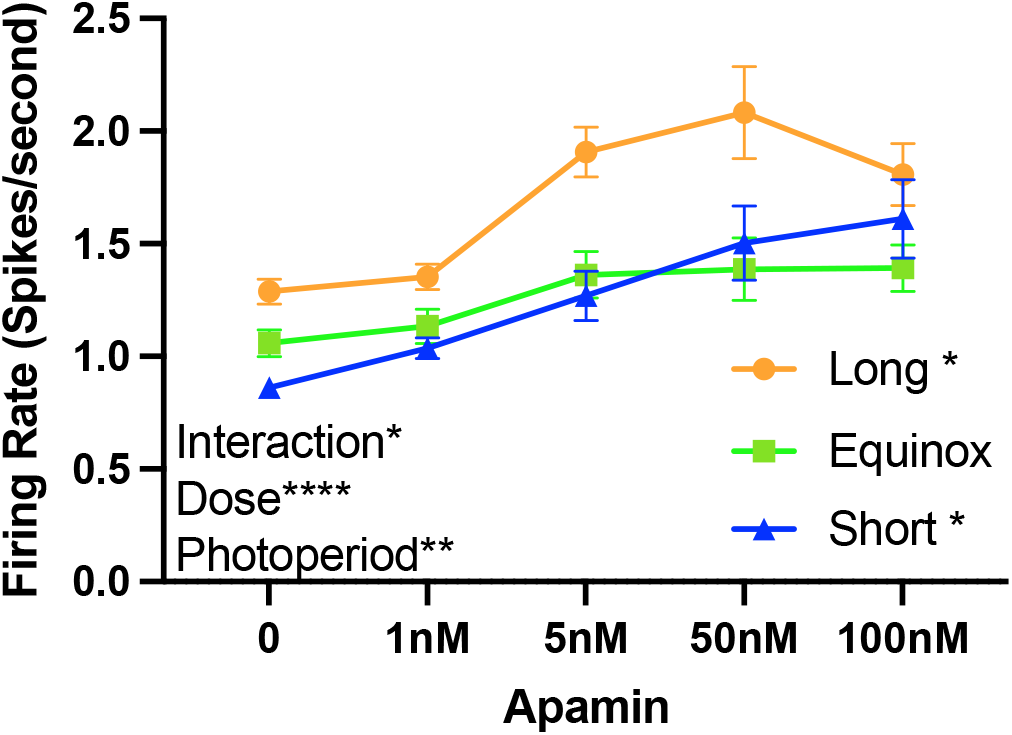
Photoperiodic Effects of Apamin on Firing Rate in Dorsal Raphe Nucleus (DRN) Serotonin Neurons. Firing rate (in spikes/second) of serotonin neurons in response to increasing concentrations of Apamin (0, 1nM, 5nM, 50nM, and 100nM) under three photoperiod conditions: Long (orange circles), Equinox (green squares), and Short (blue triangles). Repeated measures Two-way ANOVA showed significant effects for Apamin Dose (p <0.0001****), Photoperiod (p < 0.001**) and their Interaction (p <0.05*). The interaction between photoperiod and Apamin dosage shows a significant increase in firing rate at higher concentrations, with the most pronounced effect observed under short photoperiods. Tukey’s multiple comparisons revealed a significant increase in firing rate within Short and Long photoperiods in response to Apamin (adj p < 0.05*, 0 vs 100 nM).

### Reduced SK Currents in DRN Serotonergic Neurons of Long Photoperiod Mice

To examine the specific contribution of SK channel currents to the firing rate of serotonin neurons in mice exposed to different photoperiods, whole-cell patch-clamp electrophysiology was performed on neurons within the dorsal raphe nucleus. To identify serotonergic neurons, spontaneous action potentials were recorded before and after application of 8-OHDPAT (Figure 2A), as it has been previously demonstrated that serotonergic cells within the DRN selectively express 5-HT_1_A receptors, and that these receptors suppress action potential frequency (Assié & Koek, 1996; Larsson et al., 1990). Accordingly, only cells that exhibited a 50% or greater reduction in spontaneous action potential frequency in response to 8-OHDPAT were included in this study (Figure 2B). We elicited action potential and AHPs and used apamin to quantify SK-channel contributions to the AHP current under different photoperiods (Figure 2C-D). We found that the Apamin-sensitive current amplitudes collected from Long photoperiod mice were decreased compared to Equinox and Short photoperiod mice (Figure 2E) (Long mean = 14.43 pA ± 6.796; Eqx mean = 50.26 ± 9.389; Short mean = 57.06 ± 10.38; Long vs. Eqx, p = 0.0295*; Long vs. Short, p = 0.0340*; Eqx vs. Short, Tukey’s multiple comparison test: p = 0.8854). These results are consistent with our hypothesis that SK currents are reduced in dorsal raphe serotonin neurons in Long photoperiod mice consistent with the observed reduction in AHP.

**Figure 2.**
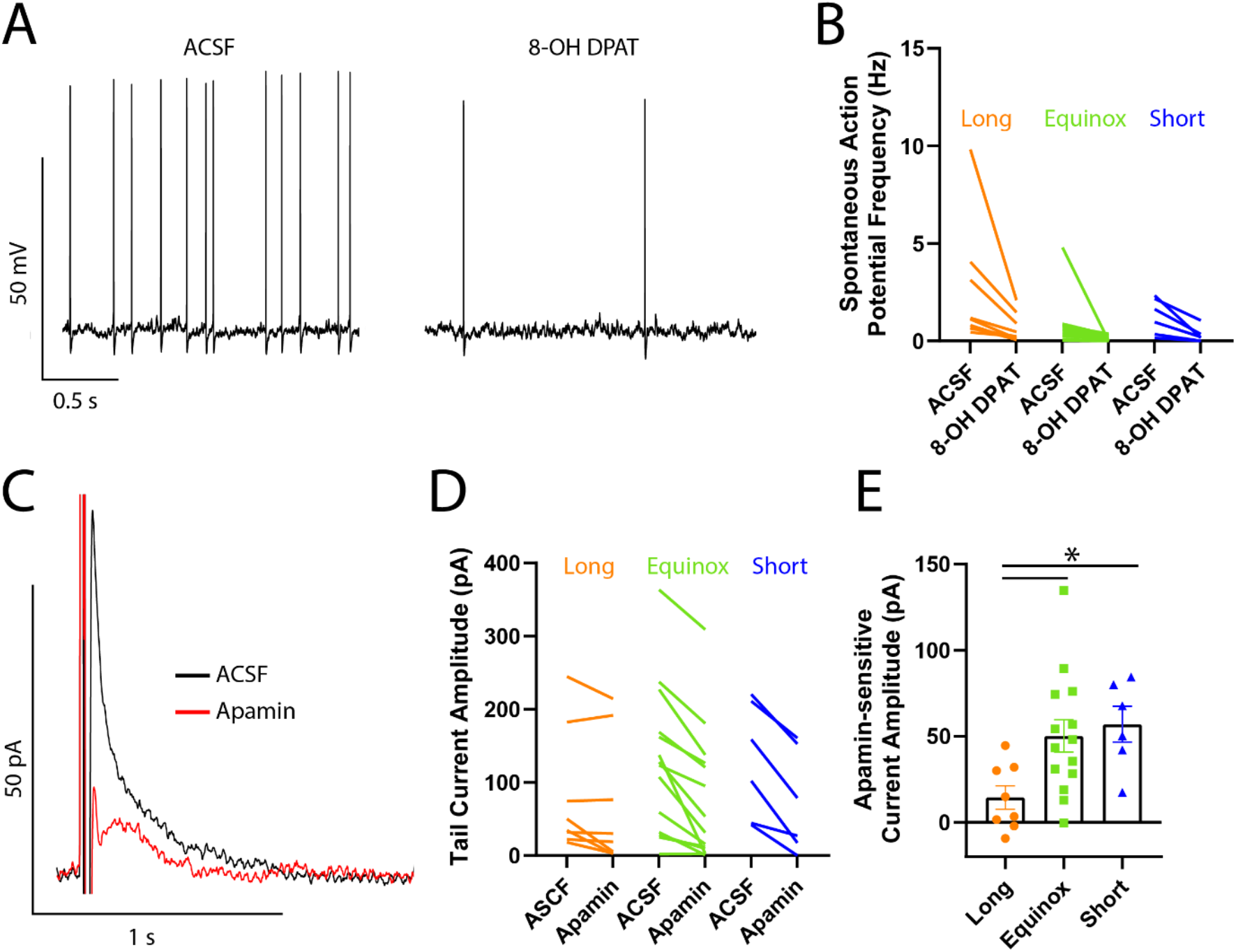
Apamin-sensitive current amplitude is reduced in Long photoperiod mice. **A)** Representative trace of spontaneous action potentials before (left) and after (right) 8-OHDPAT (5 μM) application. **B)** Summary of spontaneous action potential frequencies before and after 8-OHDPAT. Only cells which expressed a 50% or greater reduction in action potential frequency after DPAT were included in this study. **C)** Afterhyperpolarization currents were elicited by applying a 2 ms pulse to +30 mV from a holding potential of -60 mV. Representative trace of afterhyperpolarization currents before (black) and after apamin application on (red). **D)** Summary of peak amplitudes of tail currents before and after apamin wash on. Ordinary one-way ANOVA, Tukey’s multiple comparison test (Long vs. Equinox, p = 0.0295; Long vs. Short, p = 0.0340; Equinox vs. Short, p = 0.8854). Long, n = 8 cells, N = 5 animals; Equinox, n = 14 cells, N = 7 animals; Short, n = 6 cells, N = 5 animals.

### SK3 (*Kcnn3*) expression in DRN serotonin Neurons Remains Constant Across Photoperiods

The observed reduction in SK channel currents in DRN serotonin neurons could result from decreased expression of SK channels. In the dorsal raphe nucleus, the SK3 channel is the most highly expressed member of its family (Wolfart et al., 2001). To examine whether the expression levels of *Kcnn3* (SK3) transcript were altered by environmental photoperiods, we employed RNAscope to determine the expression levels of *Kcnn3* specifically in *Tph2-* expressing serotonergic neurons within the DRN (Figure 3). The analysis found that the average expression of *Kcnn3* was similar across the three photoperiod cohorts (one-way ANOVA, F = 0.624, p = 0.53). This suggests that the influence of seasonal photoperiod on SK currents occurs at the post-transcriptional level.

**Figure 3.**
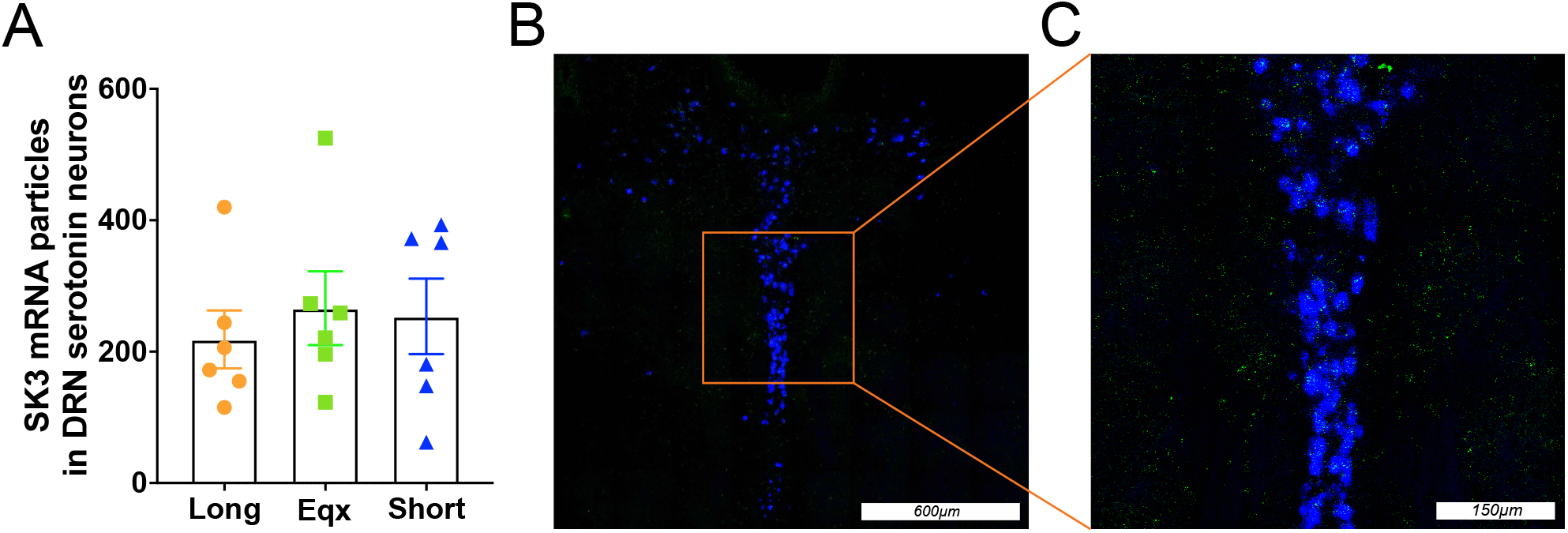
mRNA levels of *Kcnn3* (SK-3) in DRN *Tph2*-positive neurons are similar across photoperiod conditions. **A)** Quantification of colocalized *Kcnn3* and *Tph2* positive neurons in the dorsal raphe. Each bar represents the mean number of mRNA particles, with individual data points overlaid to show distribution. The error bars indicate standard error of the mean (SEM). Colocalized transcript detection of *Kcnn3* in DRN of C3Hf+/+ mice raised in Long, Equinox or Short photoperiods was similar across photoperiods (one-way ANOVA, F = 0.624, P = 0.53). **B)** Representative image of *Kcnn3* (green) and *Tph2* (blue) signal from Equinox photoperiod. **C)** Higher magnification view of serotonin neurons in the boxed region in panel B showing the distribution and colocalization of *Kcnn3* fluorescent puncta within *Tph2*-positive neurons. The brightness of *Kcnn3* signaling for panels B and C was increased by 20% in the example shown for visualization purposes.

## Discussion

The work presented here reveals that day length or photoperiod influences SK channel current amplitude in serotonin dorsal raphe neurons. Exposure to different seasonal photoperiods alters the sensitivity of serotonin dorsal raphe neurons to SK channel inhibitor apamin (Figure 1) and SK channel current amplitudes were found to be reduced in serotonin neurons of Long photoperiod mice compared to Equinox and Short photoperiod (Figure 2). Lastly, expression of *Kcnn3* (SK3) in serotonin neurons was similar across photoperiods (Figure 3), Taken together our results suggest that seasonal photoperiods act on SK channel currents responsible for the regulation of neuronal excitability, possibly at the post transcriptional level. Consistent with our hypothesis, we observe that SK currents are reduced in Long photoperiod mice compared to Short in whole cell recordings using apamin (Figure 2). This is consistent with our previous work showing that serotonin neurons from Long-photoperiod mice exhibit a lower amplitude AHP following action potentials compared to those from Short-photoperiod mice (Green et al., 2015). These changes to the electrical properties of serotonin neurons were correlated with decreased depressive and anxiety-like behaviors in Long photoperiod (Green et al., 2015). It is likely that the combined photoperiodic regulation of channels TREK-1 (Giannoni-Guzmán et al., 2020) and SK channels in DRN serotonin neurons are major drivers of these differences in behaviors across photoperiod groups.

Our results are also consistent with previous work on SK currents and social and affective behaviors. SK channel currents in dorsal raphe serotonin neurons present a reduced excitability in a chronic social isolation model, which is reversed by inhibition of SK currents using apamin (Sargin et al., 2016). Chronic isolation has been shown to increase anxiety/depression-like behaviors (Dang et al., 2015; Fone & Porkess, 2008; Koike et al., 2009; Lukkes et al., 2009; Shimizu et al., 2016; Wallace et al., 2009). Systemic administration of apamin reduced anxiety/depression-like behaviors in mice under chronic isolation (Galeotti et al., 1999; Sargin et al., 2016; van der Staay et al., 1999). These findings underscore the critical role of SK channels as dynamic modulators of serotonin neuron excitability, enabling neurons to rapidly adjust their firing properties in response to environmental cues such as photoperiod changes and social stress. By fine-tuning afterhyperpolarization, SK channels serve as a key mechanism through which external factors can shape mood and anxiety-related behaviors. Based on our previous finding that TREK-1 gene expression was different between photoperiods (Giannoni-Guzmán et al., 2020), we had hypothesized the possibility of SK channel expression in these neurons to be under photoperiodic control (Green et al., 2015). This hypothesis was consistent with work showing that SK3 null mice show enhanced hippocampal serotonin release, and antidepressant like behavioral phenotypes (Jacobsen et al., 2008). The unchanged transcript levels of SK3 (*Kcnn3*) in *Tph2* positive neurons in the DRN suggests that gene transcription and mRNA stability are not the primary factors driving the differences in channel currents observed between photoperiod conditions (Figure 3). Instead, the modulation likely occurs via post-translational mechanisms that alter channel properties without changing mRNA abundance, such as altered phosphorylation, trafficking, or subunit composition. Altered phosphorylation, mediated by kinases such as protein kinase A (PKA) or casein kinase 2 (CK2), can reduce the Ca^2+^ sensitivity of SK channels (for example, by phosphorylating the calmodulin subunit), thereby potentiating or diminishing their currents (Authement et al., 2018; Louise Faber, 2009). Altered surface expression of SK channels, whether via changes in channel endocytosis or exocytosis, can affect the density of functional channels at the plasma membrane. While direct studies in DRN are limited, similar mechanisms have been described in other brain regions where SK channel trafficking is regulated by phosphorylation-dependent interactions with scaffolding proteins (Authement et al., 2018; Louise Faber, 2009).

Our findings open several avenues for further investigation into the molecular and functional aspects of SK channel regulation by photoperiod in serotonin neurons. Future studies should aim to pinpoint the precise post-translational mechanisms—such as phosphorylation events, trafficking dynamics, or changes in subunit assembly—that drive the photoperiod-dependent modulation of SK currents. In addition, exploring how these regulatory processes vary across different developmental stages may reveal critical windows during which serotonergic circuits are most sensitive to environmental cues.

The present study contributes significantly to our understanding of how environmental factors, such as seasonal changes in day length, can shape neural function at the cellular level. By demonstrating that photoperiod influences SK channel activity in serotonin neurons, these results add to the growing mechanistic framework for linking external environmental cues with intrinsic neuronal excitability and behavior. This understanding may have profound implications for developing targeted therapies for mood disorders and other conditions associated with dysregulated serotonin signaling. Interventions designed to modulate SK channel function could, in the future, offer a novel approach to alleviating symptoms of depression and anxiety, highlighting the translational potential of this research for neuropsychiatric treatments.

## Author Contributions

M.A.G.G., J.Z, A.K. and V.M. performed the research. D.G.M., M.A.G.G., B.G, P.L. and designed the research study. P.L. and B.G. contributed essential reagents or tools. M.A.G.G. and J.Z. analyzed the data. M.A.G.G, J.Z. and D.G.M. wrote the paper.

## Acknowledgements

This research was supported by National Institutes of Health Grants NIH R01 MH108562 to DGM and the Postdoctoral program in Functional Neurogenomics grant 5T32MH065215-13.

## Data Availability Statement

The data that support the findings of this study are available from the corresponding author upon reasonable request.

## References

Andersen, S. L. (2015). Exposure to early adversity: Points of cross-species translation that can lead to improved understanding of depression. Development and Psychopathology, 27(2), 477–491. 10.1017/S0954579415000103

Assié, M.-B., & Koek, W. (1996). Possible in vivo 5-HT reuptake blocking properties of 8-OH-DPAT assessed by measuring hippocampal extracellular 5-HT using microdialysis in rats. British Journal of Pharmacology, 119(5), 845–850. 10.1111/j.1476-5381.1996.tb15749.x

Authement, M. E., Langlois, L. D., Shepard, R. D., Browne, C. A., Lucki, I., Kassis, H., & Nugent, F. S. (2018). A role for corticotropin-releasing factor signaling in the lateral habenula and its modulation by early-life stress. Science Signaling, 11(520), eaan6480. 10.1126/scisignal.aan6480

Baba, K., Pozdeyev, N., Mazzoni, F., Contreras-Alcantara, S., Liu, C., Kasamatsu, M., Martinez-Merlos, T., Strettoi, E., Iuvone, P. M., & Tosini, G. (2009). Melatonin modulates visual function and cell viability in the mouse retina via the MT1 melatonin receptor. Proceedings of the National Academy of Sciences, 106(35), 15043–15048. 10.1073/pnas.0904400106

Boulle, F., Pawluski, J. L., Homberg, J. R., Machiels, B., Kroeze, Y., Kumar, N., Steinbusch, H. W. M., Kenis, G., & Van den Hove, D. L. A. (2016). Prenatal stress and early-life exposure to fluoxetine have enduring effects on anxiety and hippocampal BDNF gene expression in adult male offspring. Developmental Psychobiology, 58(4), 427–438. 10.1002/dev.21385

Ciarleglio, C. M., Axley, J. C., Strauss, B. R., Gamble, K. L., & McMahon, D. G. (2011). Perinatal photoperiod imprints the circadian clock. Nature Neuroscience, 14(1), 25–27. 10.1038/nn.2699

Cox, O. H., Giannoni-Guzmán, M. A., Cartailler, J.-P., Cottam, M. A., & McMahon, D. G. (2024). Transcriptomic Plasticity of the Circadian Clock in Response to Photoperiod: A Study in Male Melatonin-Competent Mice. Journal of Biological Rhythms, 39(5), 423–439. 10.1177/07487304241265439

Crespi, F. (2009). Apamin increases 5-HT cell firing in raphe dorsalis and extracellular 5-HT levels in amygdala: A concomitant in vivo study in anesthetized rats. Brain Research, 1281, 35–46. 10.1016/j.brainres.2009.05.021

Damulewicz, M., Tyszka, A., & Pyza, E. (2022). Light exposure during development affects physiology of adults in Drosophila melanogaster. Frontiers in Physiology, 13. 10.3389/fphys.2022.1008154

Dang, Y., Liu, P., Ma, R., Chu, Z., Liu, Y., Wang, J., Ma, X., & Gao, C. (2015). HINT1 is involved in the behavioral abnormalities induced by social isolation rearing. Neuroscience Letters, 607, 40–45. 10.1016/j.neulet.2015.08.026

Devore, E. E., Chang, S. C., Okereke, O. I., McMahon, D. G., & Schernhammer, E. S. (2018). Photoperiod during maternal pregnancy and lifetime depression in offspring. Journal of Psychiatric Research. 10.1016/j.jpsychires.2018.08.004

Fone, K. C. F., & Porkess, M. V. (2008). Behavioural and neurochemical effects of post-weaning social isolation in rodents—Relevance to developmental neuropsychiatric disorders. Neuroscience & Biobehavioral Reviews, 32(6), 1087–1102. 10.1016/j.neubiorev.2008.03.003

Fréal, A., Rai, D., Tas, R. P., Pan, X., Katrukha, E. A., van de Willige, D., Stucchi, R., Aher, A., Yang, C., Altelaar, A. F. M., Vocking, K., Post, J. A., Harterink, M., Kapitein, L. C., Akhmanova, A., & Hoogenraad, C. C. (2019). Feedback-Driven Assembly of the Axon Initial Segment. Neuron, 104(2), 305–321.e8. 10.1016/j.neuron.2019.07.029

Galeotti, N., Ghelardini, C., Caldari, B., & Bartolini, A. (1999). Effect of potassium channel modulators in mouse forced swimming test. British Journal of Pharmacology, 126(7), 1653–1659. 10.1038/sj.bjp.0702467

Giannoni-Guzmán, M. A., Kamitakahara, A., Magalong, V., Levitt, P., & McMahon, D. G. (2020). Circadian photoperiod alters TREK-1 channel function and expression in dorsal raphe serotonergic neurons via melatonin receptor 1 signaling. Journal of Pineal Research, November 2020, 1–8. 10.1111/jpi.12705

Goldman, B. D. (2001). Mammalian Photoperiodic System: Formal Properties and Neuroendocrine Mechanisms of Photoperiodic Time Measurement. Journal of Biological Rhythms, 16(4), 283–301. 10.1177/074873001129001980

Gooley, J. J., Lu, J., Fischer, D., & Saper, C. B. (2003). A Broad Role for Melanopsin in Nonvisual Photoreception. Journal of Neuroscience, 23(18), 7093–7106. 10.1523/JNEUROSCI.23-18-07093.2003

Green, N. H., Jackson, C. R., Iwamoto, H., Tackenberg, M. C., & McMahon, D. G. (2015). Photoperiod Programs Dorsal Raphe Serotonergic Neurons and Affective Behaviors. Current Biology, 25(10), 1389–1394. 10.1016/j.cub.2015.03.050

Grunnet, M., Jensen, B. S., Olesen, S. P., & Klaerke, D. A. (2001). Apamin interacts with all subtypes of cloned small-conductance Ca2+-activated K+ channels. Pflugers Archiv European Journal of Physiology, 441(4), 544–550. 10.1007/s004240000447

Hornung, J.-P. (2010). CHAPTER 1.3—The Neuronatomy of the Serotonergic System. In C. P. Müller & B. L. Jacobs (Eds.), Handbook of Behavioral Neuroscience (Vol. 21, pp. 51–64). Elsevier. 10.1016/S1569-7339(10)70071-0

Jacobsen, J. P. R., Weikop, P., Hansen, H. H., Mikkelsen, J. D., Redrobe, J. P., Holst, D., Bond, C. T., Adelman, J. P., Christophersen, P., & Mirza, N. R. (2008). SK3 K+channel-deficient mice have enhanced dopamine and serotonin release and altered emotional behaviors. Genes, Brain and Behavior, 7(8), 836–848. 10.1111/j.1601-183X.2008.00416.x

Jameson, A. N., Siemann, J. K., Grueter, C. A., Grueter, B. A., & McMahon, D. G. (2024). Effects of age and sex on photoperiod modulation of nucleus accumbens monoamine content and release in adolescence and adulthood. Neurobiology of Sleep and Circadian Rhythms, 16, 100103. 10.1016/j.nbscr.2024.100103

Jameson, A. N., Siemann, J. K., Melchior, J., Calipari, E. S., McMahon, D. G., & Grueter, B. A. (2023). Photoperiod Impacts Nucleus Accumbens Dopamine Dynamics. eNeuro, 10(2). 10.1523/ENEURO.0361-22.2023

Koike, H., Ibi, D., Mizoguchi, H., Nagai, T., Nitta, A., Takuma, K., Nabeshima, T., Yoneda, Y., & Yamada, K. (2009). Behavioral abnormality and pharmacologic response in social isolation-reared mice. Behavioural Brain Research, 202(1), 114–121. 10.1016/j.bbr.2009.03.028

Kolb, B., & Whishaw, I. Q. (1998). Brain plasticity and behavior. Annual Review of Psychology, 49, 43–64. 10.1146/annurev.psych.49.1.43

Kuzmenkov, A. I., Peigneur, S., Nasburg, J. A., Mineev, K. S., Nikolaev, M. V., Pinheiro-Junior, E. L., Arseniev, A. S., Wulff, H., Tytgat, J., & Vassilevski, A. A. (2022). Apamin structure and pharmacology revisited. Frontiers in Pharmacology, 13, 977440. 10.3389/fphar.2022.977440

Larsson, L.-G., Rényi, L., Ross, S. B., Svensson, B., & Ängeby-Möller, K. (1990). Different effects on the responses of functional pre- and postsynaptic 5-HT1A receptors by repeated treatment of rats with the 5-HT1A receptor agonist 8-OH-DPAT. Neuropharmacology, 29(2), 85–91. 10.1016/0028-3908(90)90047-U

Lewis, P., Wild, U., Pillow, J. J., Foster, R. G., & Erren, T. C. (2024). A systematic review of chronobiology for neonatal care units: What we know and what we should consider. Sleep Medicine Reviews, 73, 101872. 10.1016/j.smrv.2023.101872

Louise Faber, E. S. (2009). Functions and Modulation of Neuronal SK Channels. Cell Biochemistry and Biophysics, 55(3), 127–139. 10.1007/s12013-009-9062-7

Lukkes, J. L., Watt, M. J., Lowry, C. A., & Forster, G. L. (2009). Consequences of post-weaning social isolation on anxiety behavior and related neural circuits in rodents. Frontiers in Behavioral Neuroscience, 3. 10.3389/neuro.08.018.2009

Manz, K. M., Siemann, J. K., McMahon, D. G., & Grueter, B. A. (2021). Patch-clamp and multi-electrode array electrophysiological analysis in acute mouse brain slices. STAR Protocols, 2(2), 100442. 10.1016/j.xpro.2021.100442

Meaney, M. J. (2001). Maternal care, gene expression, and the transmission of individual differences in stress reactivity across generations. Annual Review of Neuroscience, 24, 1161–1192. 10.1146/annurev.neuro.24.1.1161

Morin, L. P. (1999). Serotonin and the regulation of mammalian circadian rhythmicity. Annals of Medicine, 31(1), 12–33. 10.3109/07853899909019259

Morin, L. P. (2013). Neuroanatomy of the extended circadian rhythm system. Experimental Neurology, 243, 4–20. 10.1016/j.expneurol.2012.06.026

Nakane, Y., & Yoshimura, T. (2019). Photoperiodic Regulation of Reproduction in Vertebrates. Annual Review of Animal Biosciences, 7(Volume 7, 2019), 173–194. 10.1146/annurev-animal-020518-115216

Pyter, L. M., & Nelson, R. J. (2006). Enduring effects of photoperiod on affective behaviors in Siberian hamsters (Phodopus sungorus). Behavioral Neuroscience, 120(1), 125–134. 10.1037/0735-7044.120.1.125

Rincón-Cortés, M., & Sullivan, R. M. (2014). Early life trauma and attachment: Immediate and enduring effects on neurobehavioral and stress axis development. Frontiers in Endocrinology, 5(MAR), 1–15. 10.3389/fendo.2014.00033

Sargin, D., Oliver, D. K., & Lambe, E. K. (2016). Chronic social isolation reduces 5-HT neuronal activity via upregulated SK3 calcium-activated potassium channels. eLife, 5, e21416. 10.7554/eLife.21416

Shimizu, K., Kurosawa, N., & Seki, K. (2016). The role of the AMPA receptor and 5-HT3 receptor on aggressive behavior and depressive-like symptoms in chronic social isolation-reared mice. Physiology & Behavior, 153, 70–83. 10.1016/j.physbeh.2015.10.026

Siemann, J. K., Green, N. H., Reddy, N., & McMahon, D. G. (2019). Sequential Photoperiodic Programing of Serotonin Neurons, Signaling and Behaviors During Prenatal and Postnatal Development. 13(May), 1–13. 10.3389/fnins.2019.00459

Siemann, J. K., Williams, P., Malik, T. N., Jackson, C. R., Green, N. H., Emeson, R. B., Levitt, P., & McMahon, D. G. (2020). Photoperiodic effects on monoamine signaling and gene expression throughout development in the serotonin and dopamine systems. Scientific Reports, 10(1), 1–14. 10.1038/s41598-020-72263-5

Stocker, M. (2004). Ca2+-activated K+ channels: Molecular determinants and function of the SK family. Nature Reviews Neuroscience, 5(10), 758–770. 10.1038/nrn1516

Stocker, M., & Pedarzani, P. (2000). Differential distribution of three Ca2+-activated K2+channel subunits, SK1, SK2, and SK3, in the adult rat central nervous system. Molecular and Cellular Neurosciences, 15(5), 476–493. 10.1006/mcne.2000.0842

Talley, E. M., Solórzano, G., Lei, Q., Kim, D., & Bayliss, D. A. (2001). CNS distribution of members of the two-pore-domain (KCNK) potassium channel family. Journal of Neuroscience, 21(19), 7491–7505. 10.1523/jneurosci.21-19-07491.2001

van der Staay, F. J., Fanelli, R. J., Blokland, A., & Schmidt, B. H. (1999). Behavioral effects of apamin, a selective inhibitor of the SKCa-channel, in mice and rats. Neuroscience & Biobehavioral Reviews, 23(8), 1087–1110. 10.1016/S0149-7634(99)00043-3

Viejo-Romero, M., Whalley, H. C., Shen, X., Stolicyn, A., Smith, D. J., & Howard, D. M. (2024). An epidemiological study of season of birth, mental health, and neuroimaging in the UK Biobank. PLOS ONE, 19(5), e0300449. 10.1371/journal.pone.0300449

Wallace, D. L., Han, M.-H., Graham, D. L., Green, T. A., Vialou, V., Iñiguez, S. D., Cao, J.-L., Kirk, A., Chakravarty, S., Kumar, A., Krishnan, V., Neve, R. L., Cooper, D. C., Bolaños, C. A., Barrot, M., McClung, C. A., & Nestler, E. J. (2009). CREB regulation of nucleus accumbens excitability mediates social isolation–induced behavioral deficits. Nature Neuroscience, 12(2), 200–209. 10.1038/nn.2257

Wolfart, J., Neuhoff, H., Franz, O., & Roeper, J. (2001). Differential Expression of the Small-Conductance, Calcium-Activated Potassium Channel SK3 Is Critical for Pacemaker Control in Dopaminergic Midbrain Neurons. Journal of Neuroscience, 21(10), 3443–3456. 10.1523/JNEUROSCI.21-10-03443.2001

